# Second-Scale Neural Dynamics Shape Hormonal Outputs in Hypothalamic CRH Neurons

**DOI:** 10.1101/2025.03.18.641872

**Authors:** Hiroyuki Igarashi, Samuel Mestern, Aoi Ichiyama, Susheel Vijayraghavan, Julio Martinez-Trujillo, Rithwik Ramachandran, Wataru Inoue

## Abstract

The elevation of glucocorticoids is a hallmark of stress, arising from the integration of rapid neuronal signals into sustained hormonal outputs. A key interface for this neuroendocrine signal translation is corticotropin releasing hormone (CRH) neurons in the paraventricular nucleus of the hypothalamus (PVN), which release CRH at the median eminence (ME). We recently discovered that CRH_PVN_ neurons exhibit a characteristic shift in firing patterns from rhythmic short-bursts (low-activity state) to tonic firing (high-activity state) in response to stress. This raised a critical question: how are these distinct firing patterns are integrated into slower, sustained hormonal outputs at neuroendocrine terminals? Here, we implemented optical approaches to detect CRH release *ex vivo* and *in vivo.* Using newly developed sniffer cells for CRH, we measured CRH release at ME *ex vivo*, triggered by the distinct, *in vivo*-like firing patterns. Our results demonstrated that the primary determinant of neuroendocrine CRH release was the firing rate sustained over the timescale of seconds, with little contribution from specific firing patterns. These results collaborated with second-scale increase in firing rate triggered by stress stimuli. Additionally, we recorded the dynamics of CRH at the ME in the freely moving mice using genetically-encoded GPCR-activation based (GRAB) sensors for CRH. Foot shock stress triggered transient, time-locked increases in CRH release on the timescale of seconds. Importantly, these second-scale CRH pulses, when elicited during repeated foot shocks, were integrated over minutes to scale downstream hormone releases. Together, our data revealed critical roles of second-scale dynamics in CRH_PVN_ neuron activity for the neuroendocrine translation of stress signals.

## Introduction

A surge of circulating glucocorticoids (GC) represents a hallmark hormonal stress response that follows rapid behavioral and autonomic reactions to impending threats^1,2^. The protracted timescale of the hormonal response suggests its selectivity for threat signals that persist over longer periods. However, experimental evidence is limited on how rapid temporal fluctuations in neural signals are integrated into the slow hormonal responses.

The GC release is controlled by the hypothalamus-pituitary-adrenal (HPA) axis. The apex of the HPA axis is corticotropin releasing hormone (CRH) neurons in the paraventricular nucleus of the hypothalamus (PVN) whose firing activities drive the neuroendocrine release of CRH at the median eminence (ME)^2^. Recent studies revealed that the activities of CRH_PVN_ neurons *in vivo* in freely moving mice are surprisingly fast for their classic, slow hormonal response. For example, exposures to brief stressors, such as foot shock, triggered rapid and transient population activities of CRH_PVN_ neurons in the timescale of seconds, as shown by fiber photometry imaging of GCaMP fluorescence in CRH_PVN_ neurons^3–6^. Furthermore, we recently reported that *in vivo* single-unit spiking activities of CRH_PVN_ neurons increases, transiently for a few seconds, in response to pain-related nerve stimuli in anesthetized mice^7^. Interestingly, their stress-triggered firing increase involved a characteristic shift in firing patterns. Specifically, the baseline, low-activity state exhibited brief high-frequency (∼3 spikes at ∼200 Hz) bursts intervened by inter-burst intervals of around 1 second. Somewhat counterintuitively, these rhythmic bursts constrained the overall firing rate at low levels due to the long, inter-burst intervals. On the other hand, the stress-induced, high-activity state was driven by sustained, tonic spiking occurring at around 10 Hz. These specific firing patters raise a question if and how such rapid transitions in neural signals are integrated into slower hormonal outputs at the neuroendocrine terminals of CRH_PVN_ neurons. For example, does the high-frequency burst play special roles in CRH release, as has been proposed for fast neurotransmitter release^8^? Characterizing the dynamics of CRH release at high temporal resolution has been challenging due to its low amount^2,9,10^ and the lack of the juxtaposed postsynaptic targets at the ME for intrinsic signal recordings.

Here, we engineered ‘sniffer cells’^11^ for CRH, biosensor cells with picomolar-range sensitivity to extracellular CRH, that can be seeded onto acute brain slices. By electronically stimulating the ME with *in vivo*-like activity patterns, we found that the neuroendocrine CRH release is primarily controlled by the total number of spikes (*i.e.*, firing rate) over the time scale of seconds while the temporal structure of the high-frequency bursts had no impact beyond contributing to the overall firing rate. Additionally, we recorded CRH release from the ME *in vivo* using G-protein-coupled receptor activation-based (GRAB) sensor for CRH^12^. CRH release showed transient, time-locked increases on the second timescale in response to foot shock stimuli, in line with the *ex vivo* release dynamics. These brief pulses of CRH release cumulatively drove the release of adrenocorticotropic hormone (ACTH) and corticosterone (CORT) in the timescales of minutes. Together, our data revealed critical roles of second-scale dynamics in CRH_PVN_ neuron activity in regulating the HPA axis outputs.

## Results

### The firing rate controls Ca^2+^ dynamics of CRH_PVN_ neurons at the axon terminals and the soma

The ‘hormonal’ CRH levels results from cumulative release of CRH over a timescale of tens of seconds to minutes. Thus, it is plausible is that the firing rate sustained over certain time periods has a major control over the scale of hormonal CRH release. On the other hand, emerging data show that CRH_PVN_ neurons can fire *in vivo* with distinct brief high-frequency bursts (∼200 Hz) in addition to tonic firing^7^. This raises an open question about whether and how these burst might exert a non-liner influence on the intracellular Ca^2+^ dynamics and the peptide release^13–15^. To understand the relationship between firing activity, Ca^2+^ dynamics in the axon terminals at the ME, and hormonal CRH release, we designed stimulation protocols based on distinct, burst and tonic firing patterns of CRH_PVN_ neurons *in vivo*^7^. **Figure 1A** describes three electrical stimulation patterns that was delivered to the ME for 30 seconds: rhythmic bursts (RB, 3 spikes at 200 Hz/episode, total 90 spikes), tonic spikes at 10 Hz (10 Hz, total 300 spikes), and control tonic spikes that was rate-matched to RB (3 Hz, total 90 spikes).

**Figure 1.**
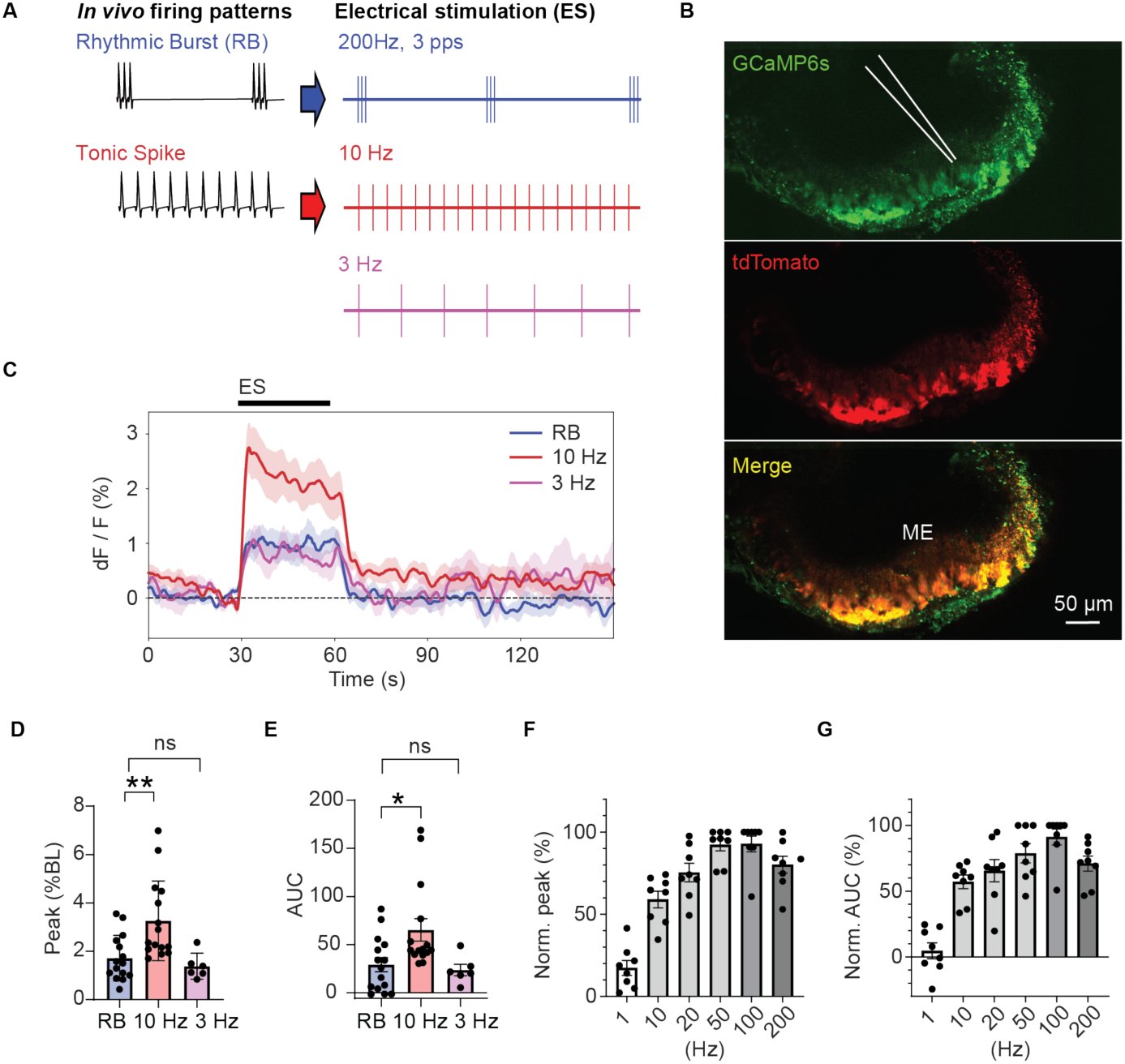
Firing rate, rather than firing patterns, controls activity-dependent Ca^2+^ elevation at the median eminence. **(A)** A schematic for *in vivo*-like the electrical stimulation (ES) protocols. **(B)** Two photon microscopy images of median eminence expressing GCaMP6s (green) and TdTomoto (red). A glass electrode (white drawing) was placed onto the ME for the delivery of the electrical stimulations. Scale bar, 50 µm. **(C)** Summary time course of GCaMP6s fluorescent signal. The bar on top depicts the duration of the ES. **(D)** The peak amplitude and **(E)** the area under the curve (AUC) of GCaMP6s fluorescent signal after electric stimulations. (RB: *n* = 15, *N* = 8, 3 Hz: *n* = 6, *N* = 3, 10 Hz: *n* = 15, *N* = 8, Kruskal–Wallis one-way ANOVA, \**p* < 0.05, \*\**p* < 0.01) **(F)** The peak amplitude and **(G)** the AUC of GCaMP6s fluorescence triggered by 5 sec ES with the range of frequency (1-200 Hz) normalized to the maximum change.

We first examined Ca^2+^ dynamics in the axon terminals of CRH_PVN_ neurons at the ME. To this end, we anterogradely expressed GCaMP6s in the axon terminals of CRH_PVN_ neurons by injecting adeno-associated virus (AAV) for cre-dependent expression of GCaMP6s (EF1α-DIO-GCaMP6s) into the PVN of CRH-Ires-Cre mice crossed with Ai14 td-tomato reporter line^16,17^. GCaMP6s were expressed in the external zone of the ME (*i.e.* an area for the hormonal CRH release), and a glass electrode was placed on the internal zone of the ME for electronic stimulation (**Fig. 1B**). The burst stimulation pattern (RB) evoked a transient increase in GCaMP fluorescence (**Fig. 1C**), indicating spike-triggered Ca^2+^ elevation in the axonal terminals. However, the magnitude of the GCaMP fluorescence increase was similar to the rate-matched tonic 3 Hz stimulation and significantly lower than the tonic 10 Hz stimulation (**Fig. 1C-E**). These results indicate that temporal structure of the high-frequency (200 Hz) bursts have little, if any, facilitatory effects on the axonal Ca^2+^ elevation, and that the total numbers of spikes (*i.e.* overall firing rate) is the primary determinant. Next, we systematically examined the effects of a range (1-200 Hz) of tonic stimulation frequencies delivered for 30 seconds: GCaMP fluorescent response displayed an increase up to 50 Hz before reaching a plateau (**Fig. 1F, G**).

To examine the somatic Ca^2+^ responses to different stimulation patterns, we repeated the same experiments in the PVN. The somatic GCaMP fluorescent increase demonstrated a similar dependence on the overall firing rate but not on the firing pattern (**Supplementary Fig. 1**). These data suggest that both at the axon terminals and somatic Ca^2+^ dynamics, which underlies the vesicular release of neurotransmitters and hormones^18^, are primarily controlled by the overall firing rate with little influence from high-frequency burst firing patterns.

### Development of CRH-sniffer cells

Detecting hormonal CRH release has been challenging due to the low signaling concentration of CRH^19^. To address this, we developed a novel cell-based biosensor for CRH, termed ‘CRH-sniffer cells’, by genetically introducing human corticotropin-releasing hormone receptor 1 (hCRHR1) tagged with mRFP, along with the cAMP Förster resonance energy transfer (FRET) probe (CEY), into HEK293 cells. CRHR1 is the class B GPCR which signals mainly through Gs coupling upon ligand binding, leading to the activation of adenylyl cyclase and subsequent increase in the intracellular cyclic AMP (cAMP)^20^. Thus, the CRH-sniffer cells report the CRHR1-induced intracellular cAMP elevation via the cAMP-dependent FRET change (**Fig. 2A**). CRH-sniffer cells with stable expression of CRHR1 and CEY were selected through drug selection, and the chosen cell line was expanded from single cell after the cell sorting based on expression levels. Confocal microscopy confirmed that the established CRH-sniffer cells uniformly expressed both CRHR1 and CEY displaying the expected cell surface expression of CRHR1, along with uniform cytoplasmic distribution of CEY (**Fig. 2B**). Analysis of a concentration-fluorescent response curve estimated an EC50 of 451 pM (**Fig. 2C**). Next, we screened a range of bioactive substances to validate the specificity of CRH-sniffer cells (**Fig. 2D**). The cells did not respond to any substances tested, except for a moderate response to noradrenaline, likely due to the intrinsic expression of adrenergic receptors in HEK293 cells^21^. CRH-sniffer cells show transient increase in response to focal pressure application (puff) of CRH and the peak response shows dose-dependent increase (Maximal Peak = 14.1%, τ_ON_ = 17.5 s for 1 µM CRH, **Supplementary Fig. 2**). The response to CRH was completely blocked in the presence of a CRHR antagonist Astressin (**Fig. 2E, F**).

**Figure 2.**
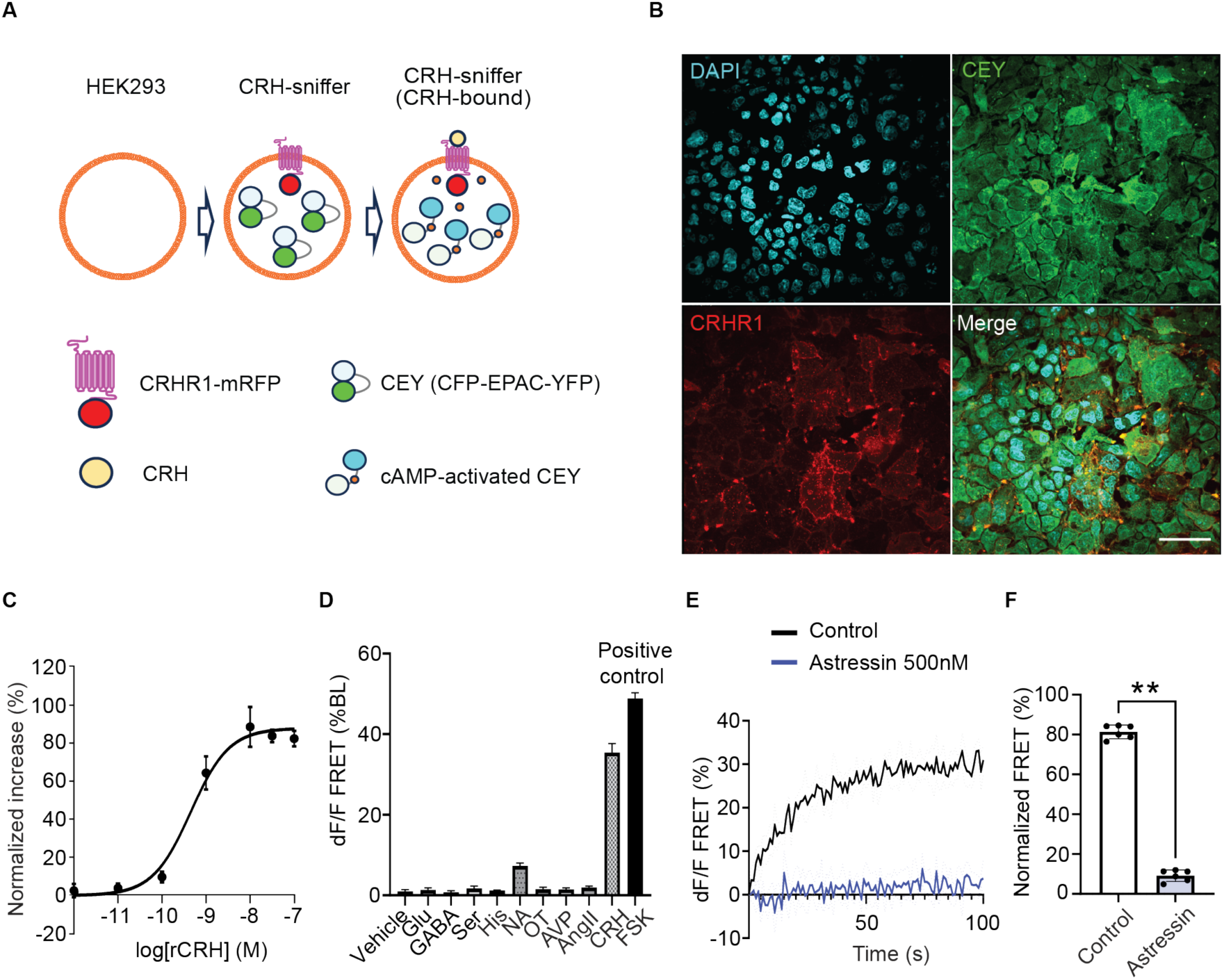
The development of CRH-sniffer cells. **(A)** Schematic illustration CRH-sniffer cells. HEK293 cells were genetically engineered to constitutively express CRHR1 and the EPAC-based sensor CEY, enabling the detection of CRH through fluorescent changes. **(B)** Confocal image of CRH-sniffer cells. DAPI, the EPAC-based cAMP sensor (CEY), hCRHR1-mRFP, and merge. Scale bar, 50 µm. **(C)** Concentration-effect curve of normalized net FRET values from CRH-sniffer cells treated with increasing recombinant CRH concentrations (n = 6). EC50 estimated at 451 pM. **(D)** CRH-sniffer cells were stimulated with various neurotransmitters and peptide hormones, and peak FRET responses within 100 s post-application were measured. Vehicle (HBSS, n = 14), Glu (1 µM, n = 7), GABA (1 µM, n = 7), Ser (1 µM, n = 8), His (1 µM, n = 8), NA (1 µM, n = 6), OT (1 µM, n = 7), AVP (1 µM, n = 7), Ang II (1 µM, n = 9), CRH (100 nM, n = 24), and FSK (20 µM, n = 25). Responses were normalized to FSK (n = number of cuvettes, 1 × 10⁵ cells each). **(E)** CRHR-antagonists Astressin (500 nM) attenuated the response to the recombinant CRH (50 nM). The pale lines represent SEM. **(F)** the bar graph shows the peak response to the 50 nM CRH with or without Astressin (*n* = 6, two-sided Mann–Whitney *U*-test, \*\**p* < 0.005).

### The firing rate controls hormonal release of CRH

To investigate hormonal CRH release at the ME, the CRH-sniffer cells were seeded onto acute brain slices containing the ME (**Fig. 3A**). Electrical stimulation of the internal zone of the ME evoked an increase in the FRET ratio in a subset of CRH-sniffer cells (**Fig. 3B**). We defined ‘responder cells’ as those with a mean dF/F (%) during the 30 seconds following the stimulation greater than the mean during the 30 seconds before stimulation. The spatial distribution of these responder sniffer cells was not confined to the vicinity of the stimulation electrode and spread along the ME (**Fig. 3B**), indicating that the electrical stimulation triggered the release of CRH from the release sites distant from the stimulation electrode. The proportion of the responders was similar across the three stimulation protocols tested (RB: 20.1 ± 1.9%, 10 Hz: 20.1 ± 1.9%, and 3 Hz: 17.9 ± 1.9%; RB vs. 10 Hz: *p* = 0.99, RB vs. 3 Hz: *p* = 0.69, one-way ANOVA with Dunnett’s multiple comparisons test). To quantify the magnitude of CRH release, we calculated the population response by summating the FRET change of all responder cells (see details in Materials and Methods). The population response triggered by the burst stimulation was similar to that of the rate-matched tonic 3 Hz stimulation and was significantly lower than that elicited by the tonic 10 Hz stimulation (**Fig. 3C, D**). These results indicate the overall firing rate, rather than the specific spiking patterns, primarily controls the magnitude of hormonal CRH release at the ME, a result in line with the Ca^2+^ response (**Fig. 1C-E**). The CRH-sniffer cell response to 10 Hz stimulation was abolished in the presence of CRH receptor antagonist Astressin (**Fig. 3E, F**), confirming the specificity of their response to CRH.

**Figure 3.**
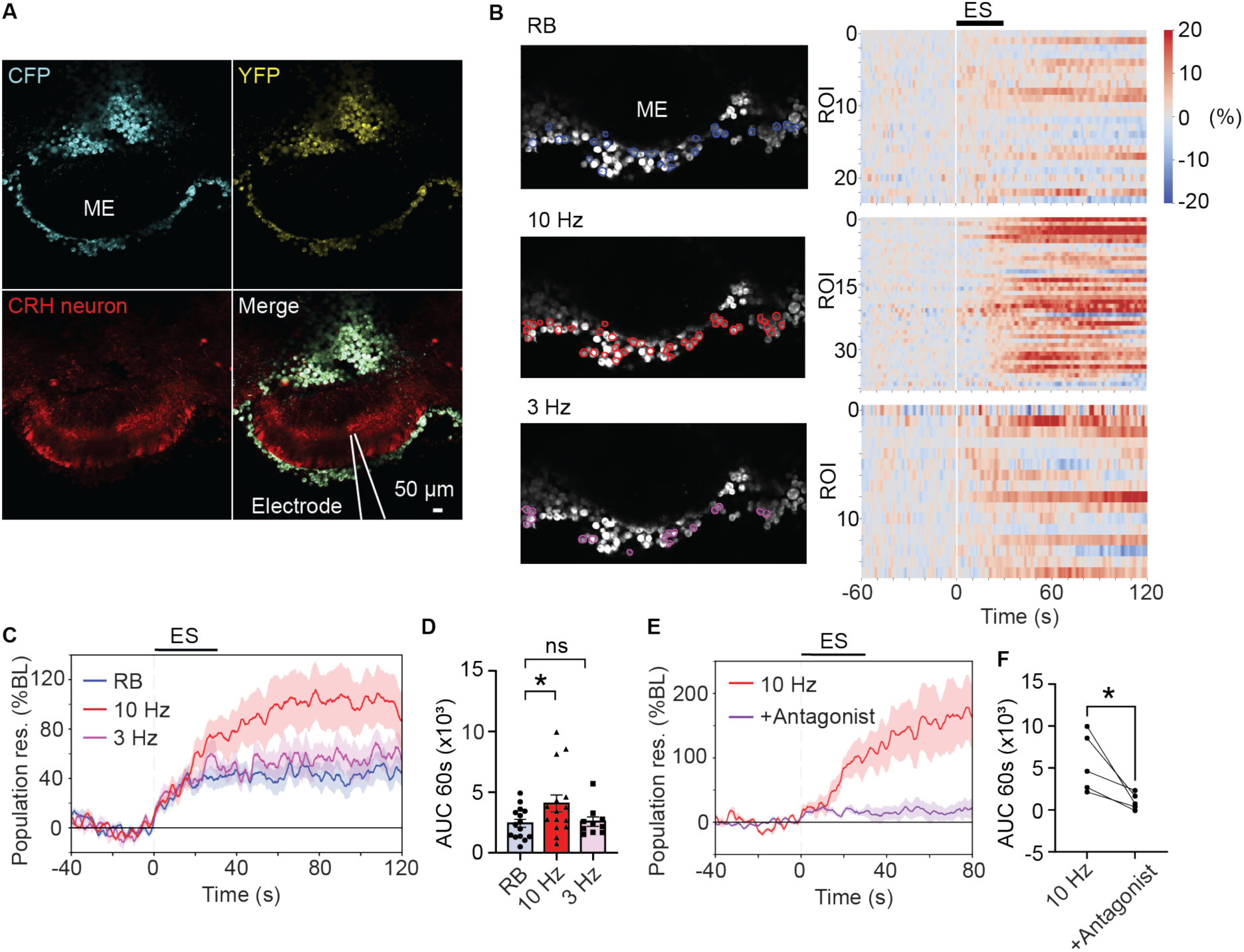
Firing rate, rather than firing patterns, controls activity-dependent CRH release at the median eminence. **(A)** Confocal image of CRH-sniffer cells seeded on top of the acute brain slices containing the ME. FRET sensor expressing CRH-sniffer cells (CFP, YFP), CRH neurons (red), and the merge are shown. **(B)** the distribution of ‘responder’ CRH-sniffer cells for each stimulation (left) and corresponding heat-map showing their response time course (right). **(C)** Summary time course for the population response of CRH-sniffer cells in response to three different stimulations patterns (RB: *n* = 15, 10 Hz: *n* = 15, 3 Hz: *n* = 10). n = number of slices **(D)** The area under the curve (AUC) during 60 sec after the ES (Mixed-effects analysis one-way ANOVA, \*\**p* < 0.005) **(E)** Summary time course for the population response of CRH-sniffer cells in response to 10 Hz stimulation in the absence and presence of CRH receptor antagonist Astressin (500 nM). **(F)** The AUC within 60 sec per cell after the SS stimulation before and after the administration of Astressin (two-tailed paired *t*-test, *n* = 5, \**p* < 0.05). Summary data are represented as mean ± s.e.m.

Besides at the ME, CRH_PVN_ neurons release CRH within and/or nearby the PVN, influencing the HPA axis activity via local feedback circuits^22–24^. We further investigated the stimulation-release relationship of the CRH in the PVN (**Supplementary Fig. 3A**). Similar to the observation in the ME, the distribution of responder cells was not restricted to the area near the stimulation electrode, indicating axonal and/or somatodendritic release (**Supplementary Fig. 3B**). The population response evoked by burst stimulation was significantly smaller than that induced by 10 Hz stimulation and similar to that of the rate-matched 3 Hz control, while the proportion of the responder cells was consistent across the three stimulation patterns (**Supplementary Fig. 3C-E**). These findings demonstrate that, in both the ME and the PVN, the overall firing rate, rather than the specific firing patterns, primarily controls CRH release.

### *In vivo* recording of CRH release at the ME in response to the pain stimuli

The foregoing *ex vivo* experiments indicated that brief spiking activities of CRH_PVN_ neurons, lasting only for a few seconds, can effectively drive firing rate-dependent CRH release. This raises a question of whether transient increases in CRH_PVN_ neuron activity *in vivo*, which can occur on the time scale of seconds^4,5^, trigger a rapid surge of CRH release in the ME. However, precise CRH release dynamics *in vivo* remain unknown, as the temporal resolutions of hormonal CRH release has been limited by microdialysis techniques, which provides cumulative release over several minutes^9^.

To address this, we investigated CRH release in the ME *in vivo* using a GRAB sensor CRF1.0^12^. We anterogradely expressed CRF1.0 in the axon terminals of PVN neurons in the ME, by injecting AAV into the PVN (AAV9-hSyn-CRF1.0). As expected, CRF1.0 expression was observed in both the PVN and ME (**Fig. 4A**). We used fiber photometry to record CRF1.0 fluorescent signals from the ME in the freely moving mice (**Fig. 4A**). Foot shock stress was used as a stimulus, as it has been shown to elicit a rapid increase in CRH_PVN_ neuron activity^4,5^. Mice were habituated to the foot shock chamber for three days, and on the fourth day, they received a single session of foot shock (2-second foot shock repeated 10 times, **Fig. 4B**). We first confirmed that this foot shock session significantly increased cFos expression in CRH_PVN_ neurons and plasma CORT levels 1 hour after stress exposure compared to the control group that did not receive foot shocks (**Supplementary Fig. 5A-C**). We found that each foot shock episode induced a time-locked, transient increase of CRF1.0 fluorescence in the ME (**Fig. 4C**). This transient increase was absent in mice expressing CRH insensitive mutant sensor (CRF_mut_)^12^, confirming the specificity of the signal (**Fig. 4C-F**). Notably, the 10 repeated shocks did not cause any sustained elevation of the signal, as indicated by the similar mean of CRF1.0 fluorescent intensity and standard deviation before and after the foot shock session (**Fig. 4H, I**). These results indicate that the repetitive, transient CRH release primarily account for the differences in the downstream hormonal response between the foot shock and control groups.

**Figure 4.**
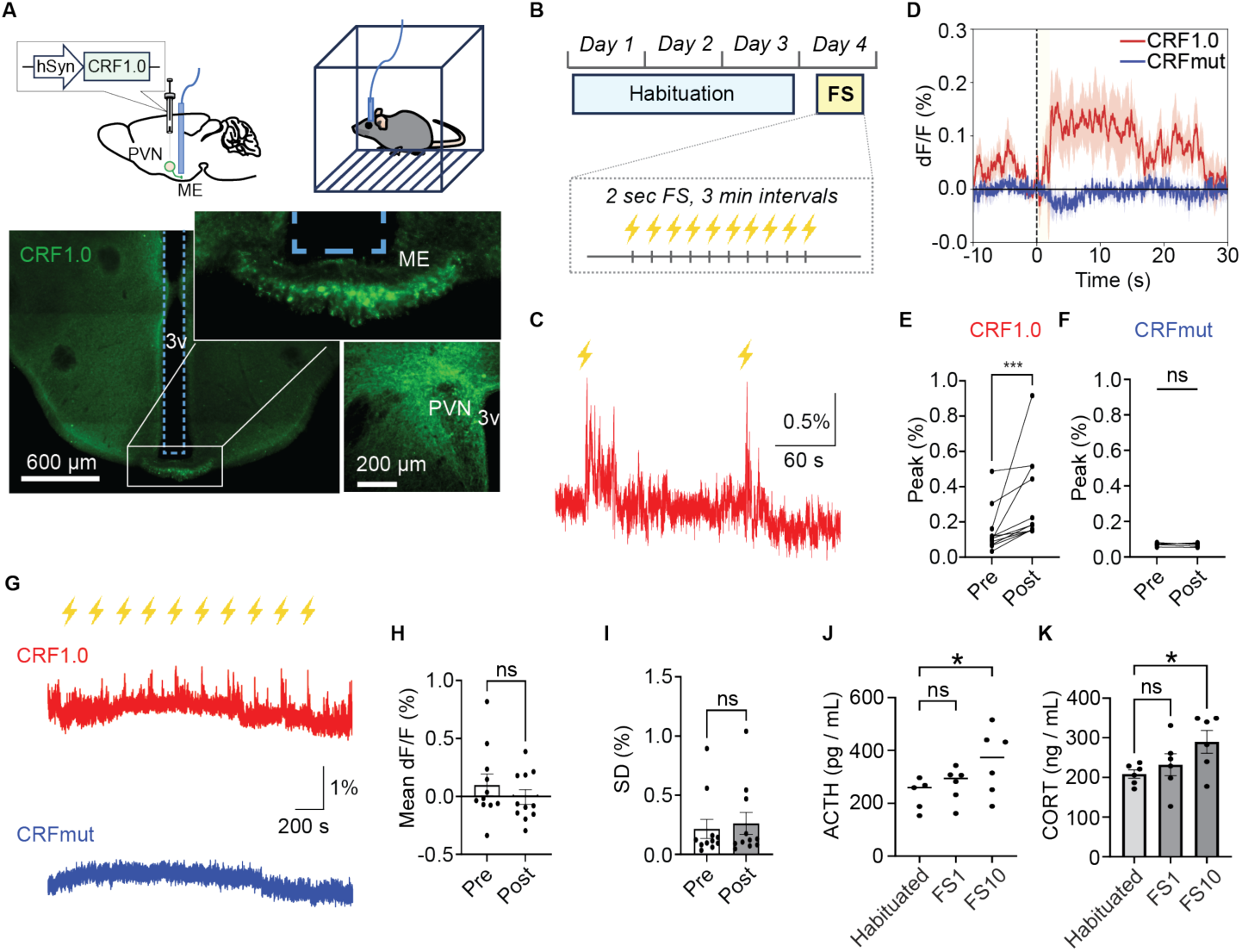
Foot shock stress elicits rapid and transient CRH release at the ME and drives a graded hormonal response *in vivo*. **(A)** A schematics for fiber photometry imaging of CRF1.0 (top). Expression of CRF1.0 (green) in the ME and the PVN, with the optic fiber placement indicated by the blue dotted line. **(B)** Experimental design for the foot shock. Mice were habituated to the behavior box for 3 days (10 min per session, twice a day), and the foot shocks (2 s/episode repeated 10 times) were given on the fourth day. **(C)** A representative trace for CRF1.0 signal in the ME in response to repeated foot shock stimuli. Yellow lightning bolts indicate the timing of foot shocks. **(D)** Summary time course of CRF1.0 (red, *N* = 11) or CRHmut (blue, *N* = 5) fluorescent signal. The dotted line indicates the timing of foot shock. **(E, F)** The peak amplitude before and after the foot shock in the mice expressing CRF1.0 (*N* = 11, \*\*\**p* < 0.005) and CRFmut (*N* = 5, ns). Wilcoxon matched-pairs signed rank test was used to compare the difference. **(G)** Representative traces of CRF1.0 (top) and CRHmut signals (middle) from the median eminence over the course of repeated foot shock (yellow lightings). **(H, I)** The mean of CRF1.0 signal (**H**) and standard deviation (SD, **I**) before and after the 10x repeated foot shock (two-tailed paired *t*-test). (**J, K**) Plasma ACTH **(J)** and CORT **(K)** levels 40 min after no foot shock (Habituated), one foot shock (FS1) and 10 foot shocks (10FS). Data were analyzed by One-Way analysis of variance (ANOVA) followed by Dunnett’s multiple comparison test (\**p* < 0.05).

Next, we hypothesized that these brief pulses of CRH release can scale the downstream hormonal response. To test this, we compared ACTH and CORT levels among three groups exposed to different numbers of foot shocks: no foot shock (FS0), a single foot shock (FS1), and 10 repeated foot shocks (FS10). Plasma ACTH levels were significantly higher in the FS10 group compared to the FS0 group, whereas the FS1 group showed similar ACTH levels to the FS0 (**Fig. 4J**). Similarly, blood CORT levels were significantly elevated in the FS10 group but not in the FS1 group compared to the FS0 (**Fig. 4K**). These data demonstrate that brief, pain-responsive pulses of CRH transiently emerge within seconds, collectively shaping the endocrine stress response.

## Discussion

The hormonal stress response follows rapid behavioral and autonomic reactions to imminent threats^1,2^. While the protracted timescale of the hormonal response indicates that rapid fluctuations of neural signals are integrated into the slow dynamics of hormone release, the mechanisms for this neuroendocrine signal integration remained poorly understood. Here, we showed: 1) CRH release at the ME is primarily controlled by firing rate over seconds, regardless of specific firing patterns of rhythmic bursts or tonic firing. 2) Stress stimuli trigger short pulses of CRH release lasting for several seconds at the ME, and these short CRH pulses are integrated over the time window of minutes to control the magnitude of downstream ACTH and CORT release. Together, our data revealed critical roles of second-scale dynamics in CRH_PVN_ neuron activity in regulating CRH release and downstream hormonal outputs.

CRH_PVN_ neurons form the apex of the HPA axis, serving as the final cellular interface that translate neural signals into hormonal CRH release. While it is widely accepted that the excitations of CRH_PVN_ neurons drives CRH release^25–28^, the precise relationship between CRH_PVN_ neurons’ spiking activity and CRH release has evaded empirical examinations. This knowledge gap was in part due to the technical difficulties in measuring inherently low amount of CRH release at the ME, which lacks the immediate postsynaptic target cells amenable for sensitive electrophysiological recordings^29^. In addition, the inquiry for this fundamental question has been hampered by the lack of information about *in vivo* spiking activities of CRH_PVN_ neurons. For the latter, we recently reported optogenetically-identified, single-unit activities of CRH_PVN_ neurons in mice and found that their activities rapidly elevates with tonic firing at around 10 Hz in response to stress stimulus (sciatic nerve stimulation mimicking pain stress)^7^. Surprisingly, the same study also found that CRH_PVN_ neurons can fire, in addition to the tonic firing, with high-frequency bursts (∼200 Hz) that are brief (∼3 spikes/burst) and intervened by long (∼1 s) inter-burst intervals. Somewhat counter-intuitively, these rhythmic brief bursts constrained the overall firing rate at low levels (*i.e.*, ∼3 Hz) due to the long inter-burst intervals and occur during non-stress conditions (and disappear in response to stress). These finding raised new questions regarding the roles of CRH_PVN_ neurons firing in CRH release. To address these questions, the present study developed CRH sniffer cells that can be seeded onto the ME in acute brain slices and detect CRH at high sensitivity in the proximity of CRH release sites. While electronic stimulation of the ME mimicking the rhythmic bursts (3 spikes at 200 Hz every second) readily triggered the release of CRH, the magnitude of CRH release was not affected by the temporal structure of the rhythmic burst and was similar to the rate-matched (3 Hz), control tonic spiking stimulation. On the other hand, tonic stimulation at 10 Hz that mimics higher, stress-associated firing rates triggered greater levels of CRH release. These results suggested that CRH_PVN_ neurons integrate spiking signals into CRH release primarily by the rate-coding. This aligns with the broader physiological framework, where the dynamics of CRH levels in the hypophyseal portal blood emerge from the cumulative release of CRH over seconds, rather than millisecond precision of release.

In addition to the classical neuroendocrine release of CRH at the ME, CRH_PVN_ neurons release CRH and transmit signals within the brain through synapses and/or volume transmission^22,23,30–33^. Recent studies by Justice, Chen, and colleagues demonstrated that optogenetic stimulation of CRH_PVN_ neurons in *ex vivo* brain slices excited nearby CRHR1-expressing neurons in a manner sensitive to CRHR1 antagonists^22,23^. These studies showed that 1 or 20 Hz optogenetic stimulation delivered over one minute caused a lasting increase in the firing rate of CRHR1-expressing neurons, suggesting that sustained tonic firing over tens of seconds triggers CRH release in the PVN. Consistent with these findings, the current study revealed that 3 and 10 Hz tonic electrical stimulation of the PVN reliably triggered CRH release after approximately ten seconds of continuous stimulation (Supplementary Fig. 3). Similar to the ME, brief bursts of high-frequency stimulation did not play any special role, and the overall firing rate controlled the magnitude of CRH release in the PVN. The importance of brief, high-frequency bursts have long been theorized for fast glutamatergic synaptic transmission, as they enhance the reliability of signal transfer to postsynaptic neurons by facilitating neurotransmitter release at synapses with low release probabilities^8,34^. Mechanistically, CRH is released via dense core vesicle fusion, whereas fast neurotransmitters are released through synaptic vesicle exocytosis^35^. It is likely that the temporal dynamics of spike trains governing these distinct release mechanisms differ, as has been observed for other peptide versus neurotransmitter release systems^36^. On the signaling side, CRH activates high affinity GPCRs in the low nM range^22,23^, whereas synaptic glutamate transmission operates via low-affinity ionotropic receptors activated in the mM range^8,34^. As such, CRHergic transmission functions on a slower temporal scale and broader spatial scale compared to traditional synaptic transmission. Taken together, rate coding over seconds, rather than brief bursts, appears more suited for regulating this type of signaling. It is worth noting that CRH_PVN_ neurons express vGlut2 and have the capacity for synaptic glutamate release outside of the PVN^27^. However, within the PVN, CRH neuron-to-CRHR1 neuron signal transmission occurs predominantly via CRH rather than glutamate^22,23^. Future studies are needed to explore the roles of burst firing in glutamatergic transmission that may occur along with CRH in growing numbers of brain areas that receive projections from CRH_PVN_ neurons^33^.

The signaling concentration of CRH is estimated in the range of 100 pM-10 nM based on CRH-stimulated cAMP elevation and ACTH release in pituitary cells^19^. To examine the extracellular CRH dynamics in this low range, we developed CRH sniffer cells that co-express Gs-coupled CRHR1 and fluorescent cAMP sensor. This design allows the sniffer cells to optically report intracellular cAMP elevation downstream of CRHR1, providing increased signal amplification. Through drug selection and cell sorting based on expression levels of CRHR1 and the cAMP sensor, we established stable and homogeneous CRH sniffer cell lines. These sniffer cells reliably report extracellular CRH levels with an EC50 of approximately 500 pM (Figure 2B), offering detection sensitivity substantially higher than currently available GRAB sensor for CRH (CRF1.0) with EC50 of 18 nM^12^. The CRH sniffer cells can be seeded onto the surface of acute brain slices for *ex vivo* optical recordings, enabling the detection of focal dynamics of extracellular CRH. The current study revealed frequency-dependent differences in CRH release at 3 Hz and 10 Hz stimulation. Additionally, in our recent study, the CRH sniffer cells revealed an attenuation of stimulus-triggered CRH release in the ME by CORT and noradrenaline^37^, further illustrating the utility of this tool for probing physiological mechanisms fine-tuning CRH release. Similar sniffer cell approaches have been used in earlier studies to detect stimulus-evoked release of oxytocin^38^ and vasopressin^39–41^ in acute brain slices. In these cases, sniffer cells co-expressed G_q/11_-coupled receptors for oxytocin or vasopressin along with genetically encoded fluorescent calcium indicators. Because sniffer cells do not require viral gene delivery, they are suitable for tissues that are less amenable to *in vivo* transfection, such as human surgical samples. However, it is important to note certain limitations of this approach. First, the ‘off’ kinetics of the CRH sniffer cells is long (>1 min at 1 nM in dissociated cells) presumably due to the combinations of ρ_off_ time constant of CRF receptors^42^ and the design of the sniffer cells reporting the intracellular cAMP elevation downstream of CRHR1 activation. Second, due to receptor desensitization^43,44^ and cAMP saturation^45^, the responsiveness of individual sniffer cells can decrease with repeated ligand exposures. In the current and recent^37^ studies, we have validated that there was no significant difference in the sniffer cell response between two repeated brain slice stimulations (3-10 Hz) with >20 min stimulation intervals. However, it is important to characterize the potential decrease in sniffer cell responsiveness in each experimental conditions when ligand exposures are repeated in the same preparations. Third, while sniffer cells have been elegantly utilized for *in vivo* measurements in earlier work^46^, their *in vivo* application remains technically challenging as it involves implantation of sniffer cells in discrete brain areas and *in vivo* two-photon imaging. For *in vivo* measurements, CRF1.0 and other genetically encoded neuropeptide sensors provides a simpler alternative, enabling fiber photometry and multiphoton microscope imaging with the added benefit of cell-type specific expression of the sensors^12^.

Using fiber photometry imaging of CRF1.0 sensor in freely moving mice, the current study revealed that stress-triggered CRH release at the ME is remarkably rapid and transient, peaking within 10 seconds and returning to baseline within 30 seconds after foot shock. This second-scale ‘pulse’ of CRH release is new and significantly advances the understanding of neuroendocrine CRH dynamics. Traditionally, the pulsatile surges of CRH in the hypophyseal portal circulation, as characterized by microdialysis, were understood to occur over the span of several minutes^9,47^. However, the real-time optical detection employed in this study unveiled a previously hidden, second-scale CRH release that was inaccessible with microdialysis due to its lower sampling frequency. Notably, this novel second-scale CRH release *in vivo* aligns well with the rapid onset of CRH release in the ME *ex vivo*, evoked by electrical stimulation. Moreover, the *in vivo* CRH release dynamics closely mirror the rapid fluctuations in the population activity of CRH_PVN_ neurons in response to foot shock^4,5^ as they displayed time-locked, transient elevation after individual shocks. Interestingly, the activity dynamics of CRH_PVN_ neurons are stressor-dependent. For instance, tail suspension elicits a transient increase in activity (<20 seconds) that is tightly time-locked to the stimulus, whereas a single intraperitoneal injection of lithium chloride, a model for visceral discomfort known to activate the HPA axis^48^, induces a prolonged elevation lasting over 20 minutes^6^. A recent study further demonstrated that CRH release in the PVN mirrors these stressor-dependent activity patterns^12^. These findings suggest that CRH release faithfully reflects the population activity of CRH_PVN_ neurons, with release kinetics adapting to the specific type of stressor and the corresponding temporal scale of CRH_PVN_ neuron activity.

Notably, repeated foot shock challenges did not lead to a persistent elevation of CRH release at the ME beyond the transient, pulsatile surges. However, these repetitive CRH pulses, even in the absence of prolonged release, significantly elevated plasma ACTH and CORT levels compared to the response elicited by a single CRH surge following a single foot shock. Thus, our data indicate that these brief pulses, occurring on a time scale of seconds, are temporally integrated over a longer time frame, such as minutes, leading to progressively amplified HPA axis activation. This mechanism may allow the system to filter out transient, inconsequential stressors while scale the hormonal response to sustained or repeated stress. In summary, our data point to a basic rule for neuroendocrine signal integration where the temporal integrations of rapid fluctuations of signals provides cumulative information to scale the protracted downstream hormonal response.

## Materials and Methods

### Animals

All experimental procedures were approved by the University of Western Ontario Animal Use Subcommittee and University Council on Animal Care in accordance with the Canadian Council on Animal Care guidelines (AUP# 2022-074). Homozygous *crh-IRES-Cre* (Stock No: 012704, the Jackson Laboratory) and *Ai14* (Stock No: 007908, the Jackson Laboratory) mice were mated, and the resulting heterozygous crh-IRES-Cre;Ai14 offspring were used as CRH-reporter mice as characterized in detail previously ^16,17,49^. All animals used were adult (>60 days old) males. They were group housed (2-4 mice per cage) in a standard shoebox mouse cage supplied with a plastic housing, paper nesting materials and wood chip bedding. The mice were housed on a 12/12 hour light/dark cycle (lights on 07:00) in a temperature-controlled (23 ± 1°C) room with free access to food and water.

### Molecular cloning and constructs

The plasmid encoding hCRHR1-mRFP was cloned by inserting the mRFP fragment into CRHR1 (wild type) cloned into pcDNA3.1+ (Invitrogen) at EcoRI (5’) and XhoI (3’) (cDNA Resource Center, Bloomsberg, PA, USA, Cat. number: #CRHR100000) using XhoI (5’) and XbaI (3’) digestion and ligation. and validated as described previously All constructs were verified by Sanger sequencing (Robarts DNA Sequencing Facility, University of Western Ontario). pCAGGS-mTurq2-EPAC-citrine (CEY)^50^ was provided by Dr. K. Okamoto (University of Toronto).

### Cell lines and culture conditions

All media and cell culture reagents were purchased from Thermo Fisher Scientific. HEK-293 cells (ATCC; Manassas, VA) were maintained in Dulbecco’s modified Eagle’s medium supplemented with 10% fetal bovine serum, 1% sodium pyruvate, and 1% streptomycin/penicillin. These chemicals were purchased from Thermo Fisher Scientific (Hampton, NH), or BioShop Canada, Inc. (Burlington, Ontario, Canada). Cells stably transfected with CMV-hCRHR1-mRFP and CAGGS-mTurq2-EPAC-citrine vectors were routinely cultured in the above media supplemented with 600 μg/ml G418 sulfate (Geneticin; Thermo Fisher Scientific, Waltham, MA).

### cAMP Signaling

Agonist-stimulated cAMP signaling was recorded in CRH-sniffer cells on a PTI spectrophotometer (Photon Technology International, Birmingham, NJ). Cells were detached in enzyme-free cell dissociation buffer, centrifuged to form a pellet (1000 rpm, 5 min), and resuspended in HBSS (Thermo Fisher Scientific). The intracellular fluorescence (excitation 434 nm; emission recorded at 474 and 529 nm) was monitored before and after addition of Glutamic acid (1 µM, G-1626, Sigma-Aldrich), GABA (1 µM, A5835, Sigma-Aldrich), Serotonin (1 µM, Sigma-Aldrich), Histamine (1 µM, Cat. No. 3545, Tocris Bioscience), Noradrenaline (1 µM, Sigma-Aldrich), Oxytocin (1 µM, Cat. No. 1910, Tocris Bioscience), Vasopressin (1 µM, Cat. No. 2935, Tocris Bioscience), Angiotensin II (1 µM, Cat. No. 1158, Tocris Bioscience), CRH (100 nM, Cat. No. 1151, Tocris Bioscience)). When tested for antagonist, cells were pre-incubated in the HBSS containing Astressin (500 nM, Cat. No. 1606, Tocris Bioscience) for 15 min then the response to CRH (50 nM) was recorded. Responses were normalized to the fluorescence obtained with Forskolin (A23187, 20 μM; Sigma-Aldrich).

The peak FRET response within 100 seconds after the drug application was measured: Vehicle (HBSS, *n* = 14), Glutamate (Glu, *n* = 7), Gamma-aminobutyric acid (GABA, *n* = 7), serotonin (Ser, *n* = 8), histamine (His, *n* = 8), noradrenaline (NA, n = 6), oxytocin (OT, *n* = 7), vasopressin (AVP, *n* = 7), Angiotensin II (Ang II, *n* = 9) were applied at 1 µM. CRH (*n* = 24) was at 100 nM while Forskolin (FSK, *n* = 25) was applied at 20 µM (n represents the number of cuvettes containing 1 × 10^5^ cells). The peak values were normalized to the FSK response.

### Stereotaxic surgery

Mice were anesthetized with 2% isoflurane using a low-flow gas anesthesia system (Kent Scientific Corporation) and placed in a stereotaxic frame. Body temperature was maintained at approximately 37°C using a heat pad controlled by a rectal temperature probe via feedback. AAV injections were performed using a Nanoject III (Drummond Scientific Company) with a pulled glass pipette filled with AAV. The pipette was lowered through a burr hole to target the PVN on each side of the brain (coordinates: A/P: −0.70 mm, M/L: ±0.25 mm, D/V: −4.75 mm from bregma). GCaMP6s expression in CRH_PVN_ neurons was achieved by injecting AAV2/9-EF1α-DIO-GCaMP6s-P2A-nls-dTomato (4 × 10^13^ GC/ml, total 800 nL; Neurophotonics Centre Viral Core, Laval University). To express CRF1.0 or the CRH-insensitive mutant CRFmut, AAV9-hSyn-CRF1.0 or AAV9-hSyn-CRFmut (2.64 × 10^13^ v.g./ml and 2.17 × 10^13^ v.g./ml respectively, total 800 nL; YL012001-AV9, YL012004-AV9, WZ Biosciences) were used. After injection, the pipette was held in place for 5 minutes to allow diffusion before being slowly retracted. For *in vivo* fiber photometry, a fiber optic cannula (400 μm core, 0.48 NA; Doric Lenses) was implanted just above the ME (coordinates: A/P: −2.10 mm, M/L: ±0 mm, D/V: −5.60 mm from bregma) and secured using adhesive dental cement. After surgery, animals were injected with analgesic (buprenorphine, 0.1 mg/kg, s.c.) and allowed to recover for at least three weeks to ensure optimal AAV expression before slice preparation.

### Acute brain slice preparation

For the slice imaging experiments, mice were deeply anesthetized with isoflurane and decapitated. Brains were then quickly removed from the skull and placed in icy slicing solution containing (in mM): 87 NaCl, 2.5 KCl, 25 NaHCO_3_, 0.5 CaCl_2_, 7 MgCl_2_, 1.25 NaH_2_PO_4_, 25 glucose and 75 sucrose (Osmolarity: 315-320 mOsm), saturated with 95% O_2_/5% CO_2_. 300 µm thick coronal sections containing the PVN or the ME were cut using a vibratome (VT1200 S, Leica). Sections were then placed in artificial cerebral spinal fluid (aCSF) containing (in mM): 126 NaCl, 2.5 KCl, 1.25 NaH2PO_4_, 26 NaHCO_3_, 10 glucose, 2.5 CaCl_2_ and 1.5 MgCl_2_ (Osmolarity: 295-300 mOsm), saturated in 95% O_2_/5% CO_2,_ maintained at 36°C for 30 minutes, and thereafter kept at room temperature in the same aCSF for the rest of the day.

Sniffer cells were dissociated in warm aCSF at the concentration of 10,000 cells/µl and applied to the acute brain slices. For the imaging of sniffer cells adhered to glass coverslips, the cells were reconstituted into in growth medium and plated at 5000 cells/cm^2^ on 13 mm borosilicate glass coverslips, pretreated with poly-L-lysine (Sigma, P4707). Cells were given 48 hours to grow before imaging.

### Multiphoton Imaging

Acute slices were imaged using a multiphoton microscope (Bergamo II, Thorlabs Imaging Systems, Sterling, VA) with a 20× objective lens (XLUMPLFLN, 1.0 NA, water immersion, Olympus, Tokyo, Japan). For the GCaMP recordings, excitation was at 940 nm and the emission signals were detected through 562 nm edge epi-fluorescence dichroic mirror (FF562-Di03-32×44-FX, Semrock, Rochester, NY) and 525/50 nm single-band bandpass filter (FF03-525/50-32, Semrock). mRFP and tdTomato signals were obtained through 607/70 nm single-band bandpass filter (FF01-607/700-32, Semrock). GCaMP images were acquired at 15.1 Hz for the PVN and 15.3 Hz for the ME.

The Ca^2+^ response was quantified based on the square ROI (500 µm × 200 µm) placed to enclose the ME. The peak and area under the curve (AUC) of ΔF/F were calculated within a 30-second window set on the pre (baseline) and post-ES period. For the PVN imaging data, the whole soma of each CRH neuron was segmented for analysis using cellpose^51^, and the average response was used in subsequent analyses.

For CRH-sniffer cell imaging, 800 nm laser (Chameleon, Coherent Laser Group, Santa Clara, CA) was used and the emission signals were detected through 495 nm edge epi-fluorescence dichroic mirror (FF495-Di03-25×36, Semrock, Rochester, NY) and 447/60 (FF02-447/60-25, Semrock) nm and 525/50 nm single-band bandpass filter (FF03-525/50-32, Semrock) with the identical gain for two channels. Imaging frequency was set at 15.1 Hz. The ROIs corresponding to the sniffer cells were segmented by cellpose^51^ based on the single optical plane. Unstable ROIs due to z-axis drift were removed based on a criterion for the SD (pre-recording SD < post-recording SD) and peak threshold (30% > ΔF/F > -20%). ‘Responder cells’ to ES were defined if the mean ΔF/F (%) calculated from 30 sec of post-ES period was bigger than the pre-ES 30 sec. The time course for the population response were calculated as follows: First, the time course of the response for individual sniffer cells were derived as the ΔF/F value for each time window. Next, the ΔF/F value from all responder cells within the same trial were summed to generate the time course of the population response.

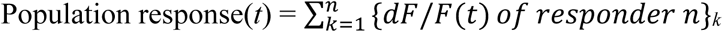

where *n* represents the total number of responder cells, and *t* takes the time point based on the sampling time.

### Fiber Photometry

The fluorescence signals were recorded with the photometry system equipped with a fluorescent mini-cube (Doric Lenses) to transmit sinusoidal 465 nm LED light modulated at 572 Hz and a 405 nm LED light modulated at 209 Hz. LED power was set at 25 μW. Fluorescence was collected through the patch-cord connected to the optic fiber implant of each mouse and transmitted back to the mini-cube, amplified, and focused into an integrated high sensitivity photoreceiver (Doric Lenses). As has been done in a previous study^3^, a linear regression was used to correct for bleaching of the signal using the slope of the 405 nm signal fitted against the 465 nm signal, where

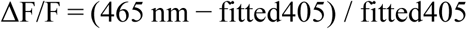

In the analysis, AUC was calculated as dF × sampling time for each time point, and AUC_30s was estimated as the integral of dF/F value over 30 seconds of the pre- and post-foot shock period. The peak takes the maximal dF (%) within 10 sec before and after each FS episode. The time window for calculating the mean, AUC, and SD was defined as follows: the ‘pre’ period is 30 seconds before the first foot shock, and the ‘post’ period spans from 30 to 60 seconds after the 10th foot shock.

### Foot Shock

Mice were habituated to the foot shock chamber and handling procedure twice daily (10 min per session, with a morning session between 9 a.m. and 12 p.m., and an afternoon session between 2 p.m. and 5 p.m.) for three consecutive days. The multi-conditioning system (TSE systems) was set at a background noise level of 65 dB and equipped with an infrared light for video visualization^52^. Then, mice were habituated for 5 min before receiving foot shock stimuli (10 trials of 2 s in length, foot shock intensity of 0.5 mA, inter-trial intervals of 1 or 3 min) and remained in the arena for 8 min after the last foot shock before being transferred to their home cage. Each foot shock was coupled to the delivery of a TTL output for synchronization with the fiber photometry recordings. On the fourth day, they received foot shock. The blood samples were collected right after the removal of the mice from the shock chamber. Control mice were habituated and handled identically without foot shocks.

### Immunohistochemistry

C-Fos expression was examined 1 hour after the foot shock experiments. The mice were deeply anaesthetized with sodium pentobarbital (100 mg/kg, intraperitoneally). A tail-pinch test was performed to ensure the depth of the anesthesia. The mice were then transcardially perfused with ice-cold saline solution (0.9% NaCl) followed by 4% paraformaldehyde (PFA, Sigma) dissolved in phosphate-buffered saline (PBS, pH 7.4). The brains were removed and fixed in 4% PFA at 4°C overnight. The brains were then coronally sectioned into 50 μm thick slices using a vibratome (VT1000S, Leica Biosystems, Concord, ON, Canada). Sections were stored in a cryoprotectant solution (30% glycerol, 30% ethylenglycol, in 20 mM PB) at ‒20°C until use. Immunohistochemistry was performed on free-floating sections. Sections were rinsed in PBS 3 times for 5 minutes each and then incubated in a blocking solution (3% normal donkey serum, 0.3% Triton X-100 and 0.03% NaN_3_ in PBS) for 1 hour. Slices were then incubated with anti-c-Fos rabbit monoclonal antibody (Cell Signalling Technology, cat: 2250S, 1:1000 dilution in blocking solution) for overnight at room temperature. After three washes in PBS, the sections were incubated with Alexa Fluor 647 donkey anti-rabbit IgG (ThermoFisher Scientific, cat: A-31573, 1:500 dilution in blocking solution). After three washes with PBS, the sections were incubated in 4’,6-diamidino-2-phenylindole (DAPI, 100 ng/ml in PBS, Sigma, cat: D9542) for 10 minutes. After two washes with PBS, the sections were mounted on glass slides and cover slipped using Fluoromount-G mounting medium (Electron Microscopy Sciences (EMS), cat: 17984-25).

c-Fos immunohistochemistry sections were imaged and analyzed as follows. Z stack images (0.685 µm thick optical sections, 20-23 sections) were obtained with a confocal microscope (Leica SP8, Leica-Microsystems) using a 20x objective (HC PL APO CS2, 0.75 NA, dry, Leica-Microsystems). Experimenters were blinded for treatments prior to image analysis. Each image was reconstructed in 3D and quantified using Imaris (Imaris v7.6.4, Bitplane, AG, Zurich, Switzerland). To estimate the volume positive for tdTomato and c-Fos expression, automatic thresholding was applied and a surface was created for each channel. The same threshold values were then used to create a colocalization channel dual-positive for tdTomato and c-Fos. A third surface was then created for the colocalization channel, determining the volume of the colocalization. The degree of cFos expression by tdTomato-positive CRH neurons was derived from colocalization volume divided by tdTomato volume for individual images.

CRF1.0 and CRFmut were detected by anti-GFP antibody (1:1000, ab13970, abcam) and goat anti-chicken IgY secondary antibody conjugated with Alexa Fluor 488 (1:500, A11039, invitrogen). The sections were imaged through LPlanFL PH2 10x objective lens equipped to the EVOS FL Auto 2 (Thermo Scientific). The fluorescence of CRF1.0 and CRFmut was excited and filtered by the EVOS LED light cube for GFP (Ex. 470/22 nm and Em. 525/50 nm, AMEP4651) and DAPI was detected by DAPI light cube (Ex. 357/44 nm and EM. 447/60 nm, AMEP4650).

### Hormone Assays

Blood samples were collected 1 hour or 40 minutes after the experiments from the same group of mice used for immunohistochemistry or the behavioral studies, respectively. Each serum sample was analyzed in duplicate with each of the two commercially available ELISA kits ELISA kits: ACTH (MDB, M046006) and CORT (Arbor Assays, K014-H5W) according to the manufacturers’ instructions.

### Statistics

All statistical analysis were performed using the GraphPad Prism 7 (Graphpad Software Inc., San Diego, CA). For c-Fos immunohistochemistry, two to four images of the PVN were obtained per animal, and the average value for an individual animal was considered as a N of 1. For *ex vivo* slice imaging, an individual slice was considered as a n of 1 for the statistical analysis. The number of animals in a treatment group as shown as N. To perform a two-group comparison, a paired or unpaired *t*-test and Mann-Whitney test were used after confirming the data sets fits to a Gaussian distribution or not. To compare multiple groups, a one-way ANOVA was performed. *p* < 0.05 was considered statistically significant.

## Acknowledgements

We thank all members of Inoue lab for thoughtful inputs to the project. We are especially grateful to Ms. Irma Meteluch for her help with animal husbandry and Mr. Jian Xiang Ding for his help with histology. We thank Dr. Kenichi Okamoto (University of Toronto) for kindly providing pCAGGS-mTurq2-EPAC-citrine (CEY) construct. We are grateful to Dr. Yulong Li and Dr. Huan Wang (Peking University) for their advice on CRF1.0 sensor. We also thank Dr. Karl Iremonger and Dr. Emmet Power (University of Otago) for their constructive feedback on our draft. Special thanks go to Dr. Miguel Skirzewski (Western University) and the Rodent Cognition RIC (BrainsCAN, Western University) for their support with the photometry system. This work was supported by the BraisCAN, JSPS fellowship, and Konica Minolta Imaging Science and Technology Foundation to HI, and BrainsCAN and the Canadian Institute of Health Research Project Grant to WI.

## Author contributions

HI designed experiments, acquired, analyzed and interpreted data, and drafted the article. SM, AI, SV and RR acquired, analyzed and interpreted data. J.C.M.-T. and RR conceived the project. WI conceived, designed experiments, interpreted data and drafted the article.

---

**Supplementary Figure 1.**
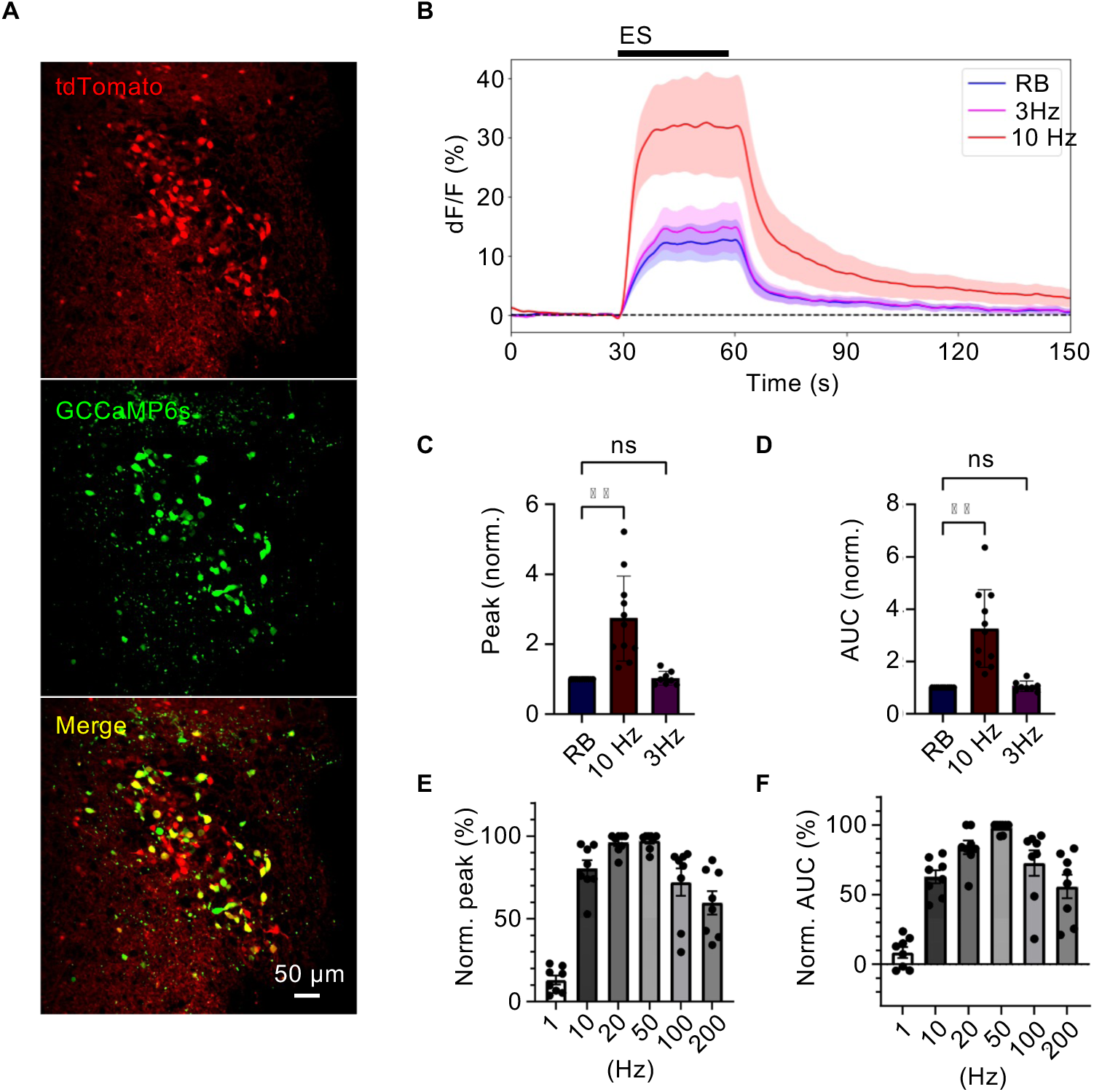
Activity-dependent calcium response in the PVN. (A) Expression of Ca^2+^ indicator GCaMP6s in the PVN. CRH neurons (red), GCaMP6s positive cells (green), and the merge (yellow) are shown. (B) Population dynamics of Ca^2+^ signal generated in response to the rhythmic burst (RB, blue), 3 Hz (magenta), or 10 Hz (red) stimulation (*n* = 7). Top grey bar represents the duration of the stimulation. The somatic response of CRH(+)/GCaMP6s(+) neurons in the PVN were analyzed. (C) Quantitative comparison of the peak amplitude calculated from the calcium responses to electronic stimulations. (D) Quantitative comparison of the area under the curve (AUC). (E, F) Frequency dependent change of peak amplitude (E) and AUC (F).

**Supplementary Figure 2.**
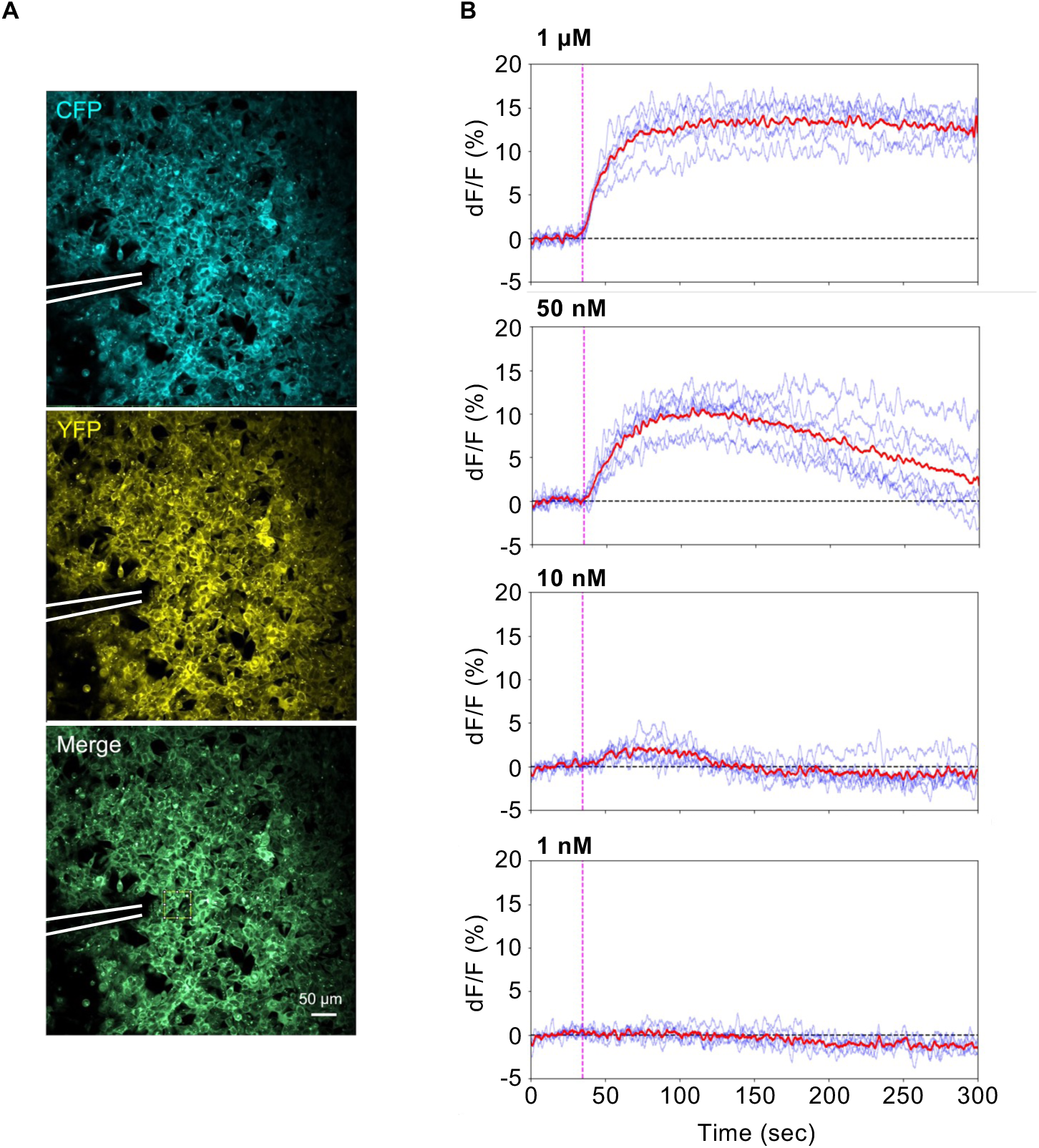
The response kinetics of CRH sniffer cells. (A) CRH sniffer cells were seeded on the coverslip, and a glass pipette for focal puff application of CRH (white lines) was positioned right above the cells. The square ROI was created to measure the response (yellow line in the bottom image). (B) Recombinant CRH was puff applied onto the CRH sniffer cells at 1 µM (*n* = 6, top), 50 nM (*n* = 7, 2^nd^ from top), 10 nM (*n* = 6, 3^rd^ from top), and 1 nM (*n* = 5, bottom). The magenta dot lines indicate the puff timing. The blue lines show individual response, and the averaged trace is shown in red.

**Supplementary Figure 3.**
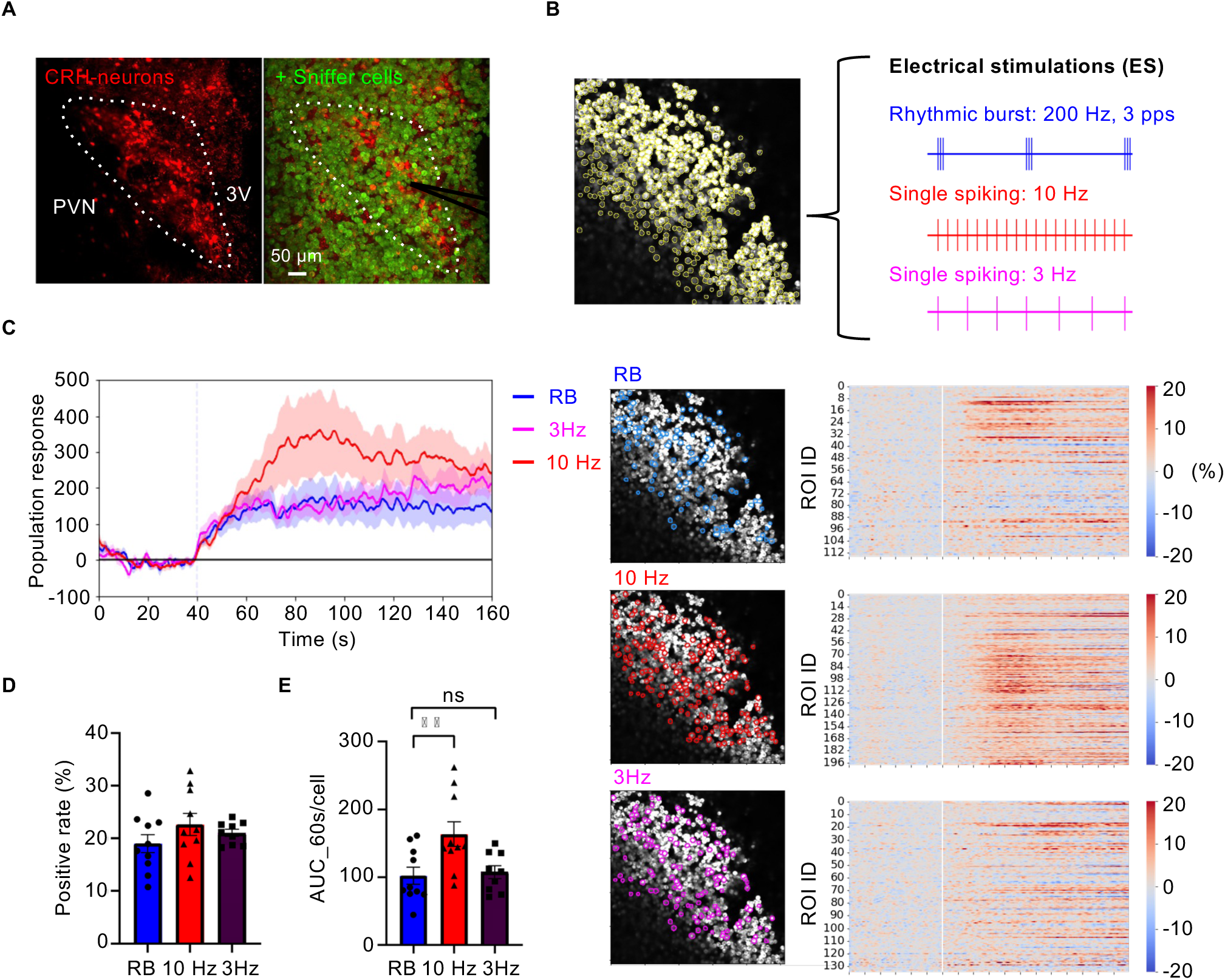
Sniffer cell response to the CRH release in the PVN. (A) Two photon microscopy image of CRH-sniffer cells seeded on top of the acute brain slice containing the PVN. The white dotted line delineates the PVN. 3V, the third ventricle. (B) The response of CRH sniffer cells against each electrical stimulation. The yellow contour shows the location of the sniffer cells detected by segmentation (top). The location of responsive sniffer cells are shown by the colored lines (blue: RB, red: 10 Hz, magenta: 3 Hz) and individual response from responded ones are shown as the heatmap on the side. (C) the populational change of AUC corresponding to the CRH release triggered by each stimulus(RB: *n* = 10, 10 Hz: *n* = 10, 3 Hz: *n* = 9). (D) the rate of positive sniffer cells emerged after the ES. (E) Individual response from a single responsive sensor cell to each ES pattern.

**Supplementary Figure 4.**
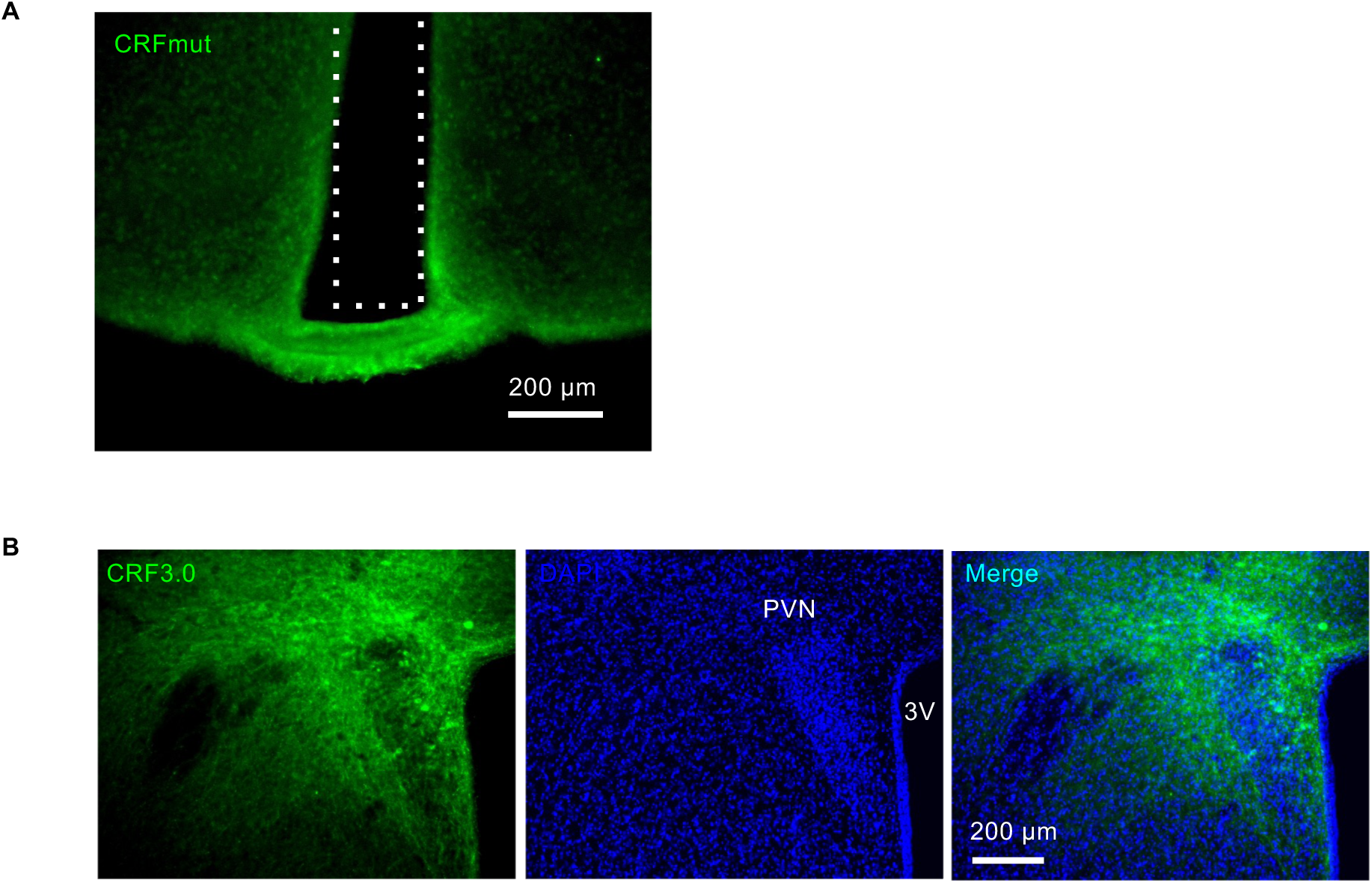
Photometry recording of CRH release at the ME. (A) The expression of CRH insensitive mutant sensor (CRFmut) at the ME. The optic fiber tract is marked with the dotted line in white. (B) The expression of CRF3.0 in the PVN (green) with the nuclear staining by DAPI (blue).

**Supplementary Figure 5.**
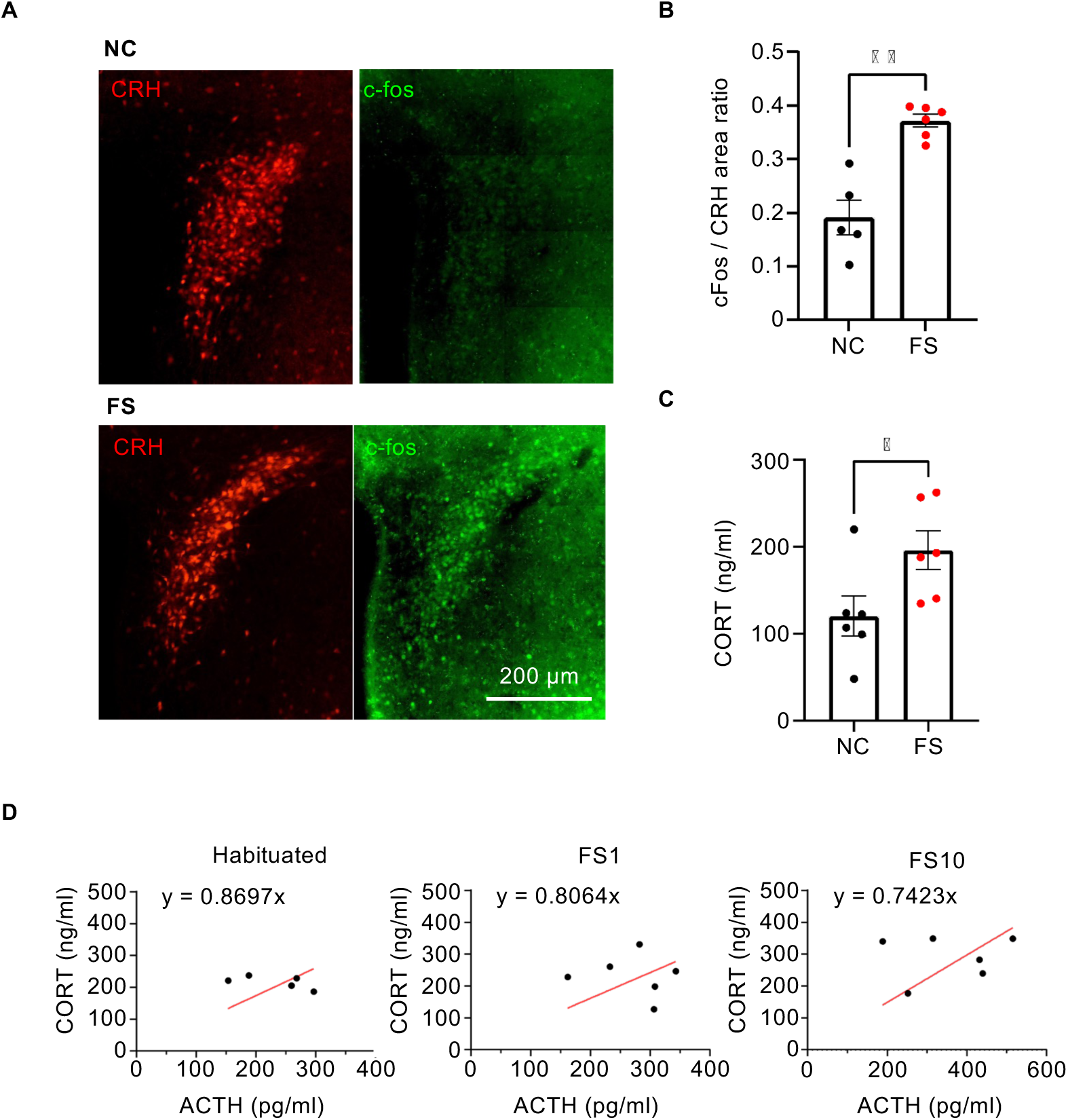
The activation of the HPA axis by foot shock stress. (A) Representative images of CRH neurons (red) and c-fos-immunoreactivity (green) in the PVN after the foot shock. Top: negative control (NC) animal which were habituated to the chamber but not given any foot shocks. Bottom: Foot shocked animal which underwent 10 times foot shocks (FS10). (B) Quantified data showing the ratio of total c-fos immunoreactive area divided by the total CRH positive area (\*\**p* < 0.005, Welch’s *t*-test). (C) Plasma corticosterone (CORT) level of both NC and FS group (\**p* < 0.05, unpaired *t*-test). (D) Correlation between ACTH and CORT in the habituated(left), FS1(middle), and FS10(right) group. The equation of the fitting curve is shown on each graph.

