## Supplemental Figures for "Second-Scale Neural Dynamics Shape Hormonal Outputs in Hypothalamic CRH Neurons"

Supplementary Figure 1

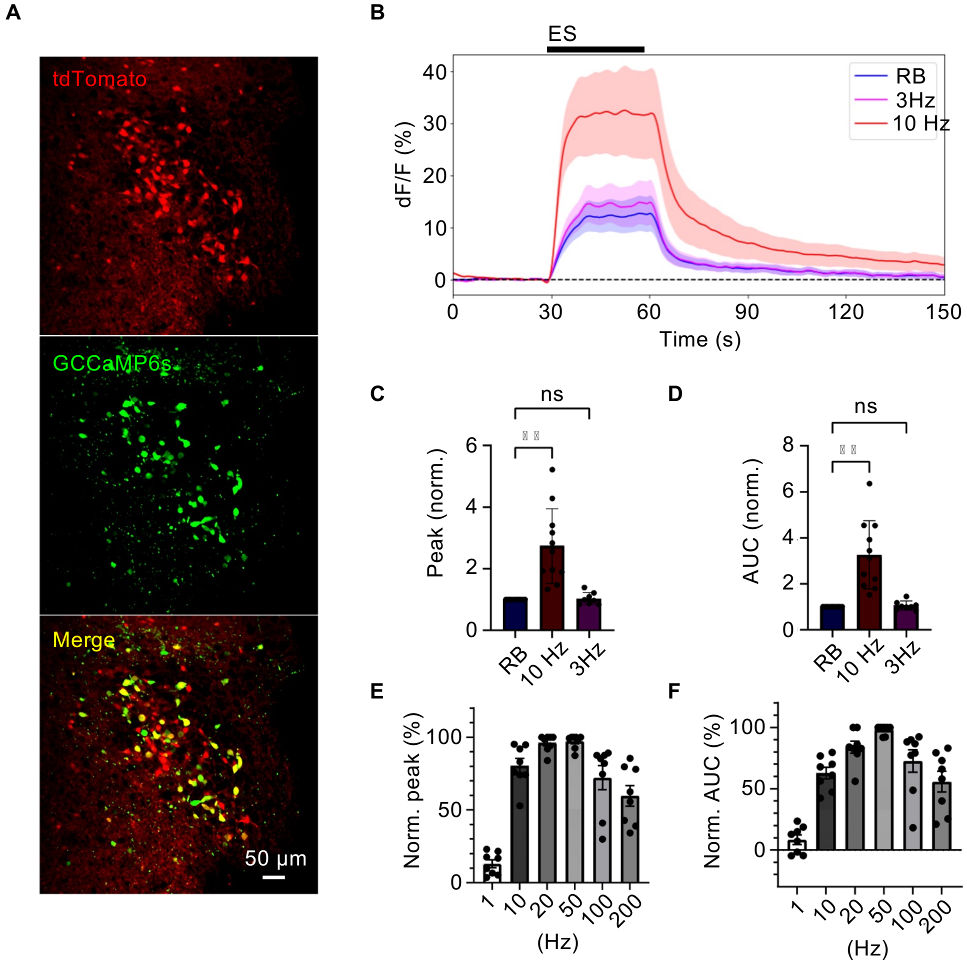


**Supplementary Figure 1. Activity-dependent calcium response in the PVN.**
(A) Expression of $Ca^{2+}$ indicator GCaMP6s in the PVN. CRH neurons (red), GCaMP6s positive cells (green), and the merge (yellow) are shown. (B) Population dynamics of $Ca^{2+}$ signal generated in response to the rhythmic burst (RB, blue), 3 Hz (magenta), or 10 Hz ( red) stimulation ($n$ = 7). Top grey bar represents the duration of the stimulation. The somatic response of CRH(+)/GCaMP6s(+) neurons in the PVN were analyzed. (C) Quantitative comparison of the peak amplitude calculated from the calcium responses to electronic stimulations. (D) Quantitative comparison of the area under the curve (AUC). (E, F) Frequency dependent change of peak amplitude (E) and AUC (F).

Supplementary Figure 2

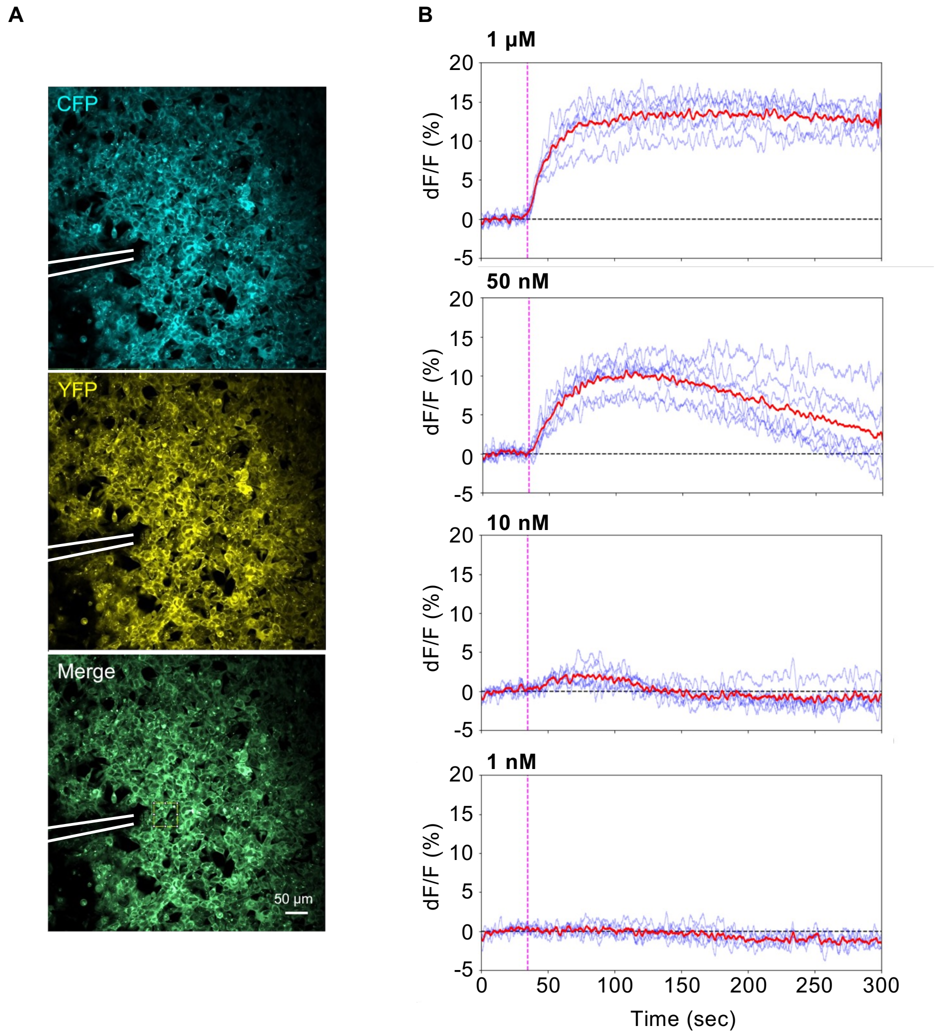


**Supplementary Figure 2. The response kinetics of CRH sniffer cells.**
(A) CRH sniffer cells were seeded on the coverslip, and a glass pipette for focal puff application of CRH (white lines) was positioned right above the cells. The square ROI was created to measure the response (yellow line in the bottom image). (B) Recombinant CRH was puff applied onto the CRH sniffer cells at 1 µM (*n* = 6, top), 50 nM (*n* = 7, 2nd from top), 10 nM (*n* = 6, 3rd from top), and 1 nM (*n* = 5, bottom). The magenta dot lines indicate the puff timing. The blue lines show individual response, and the averaged trace is shown in red.

Supplementary Figure 3

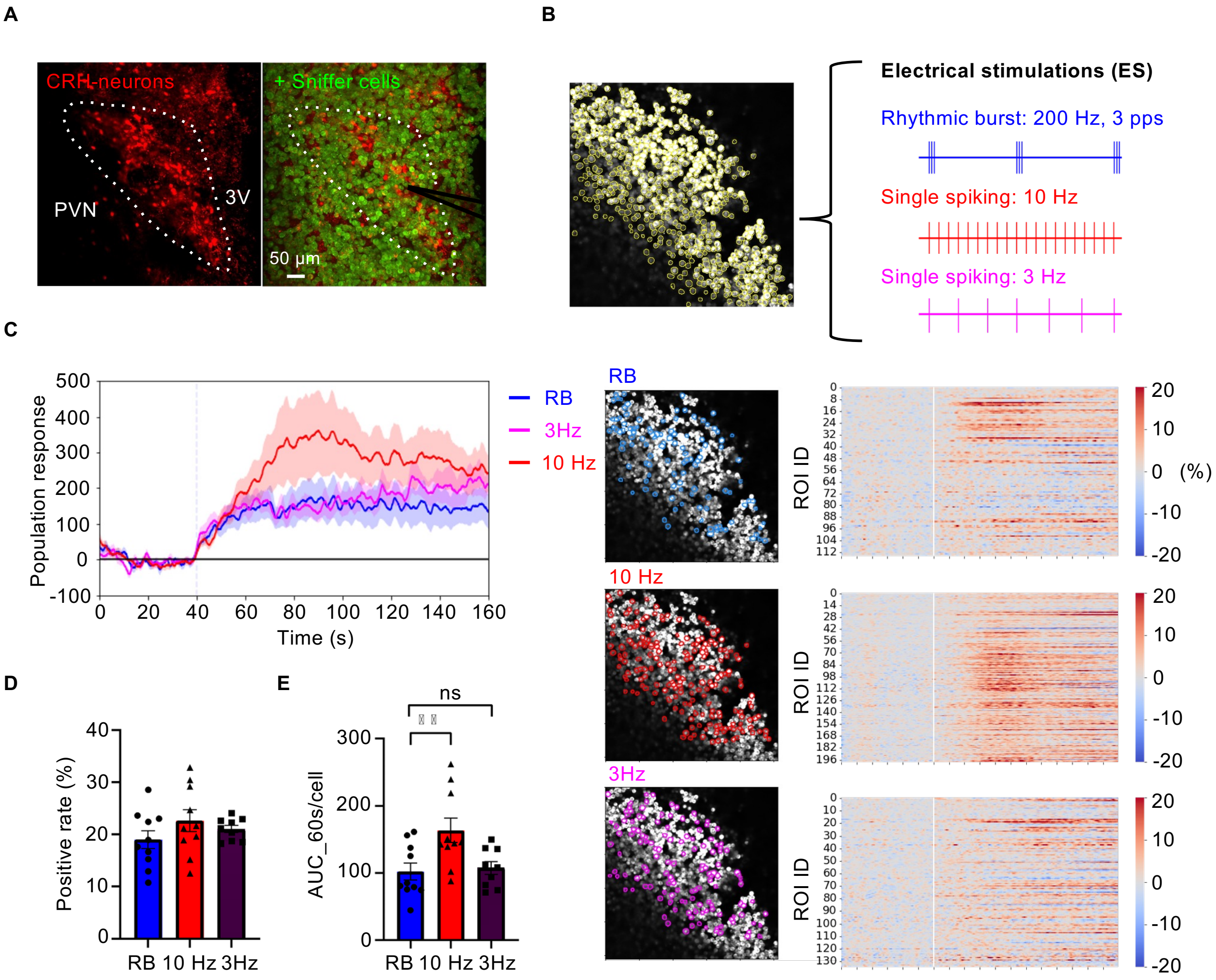


**Supplementary Figure 3. Sniffer cell response to the CRH release in the PVN.**
(A) Two photon microscopy image of CRH-sniffer cells seeded on top of the acute brain slice containing the PVN. The white dotted line delineates the PVN. 3V, the third ventricle. (B) The response of CRH sniffer cells against each electrical stimulation. The yellow contour shows the location of the sniffer cells detected by segmentation (top). The location of responsive sniffer cells are shown by the colored lines (blue: RB, red: 10 Hz, magenta: 3 Hz) and individual response from responded ones are shown as the heatmap on the side. (C) the populational change of AUC corresponding to the CRH release triggered by each stimulus(RB: *n* = 10, 10 Hz: *n* = 10, 3 Hz: *n* = 9). (D) the rate of positive sniffer cells emerged after the ES. (E) Individual response from a single responsive sensor cell to each ES pattern.

Supplementary Figure 4

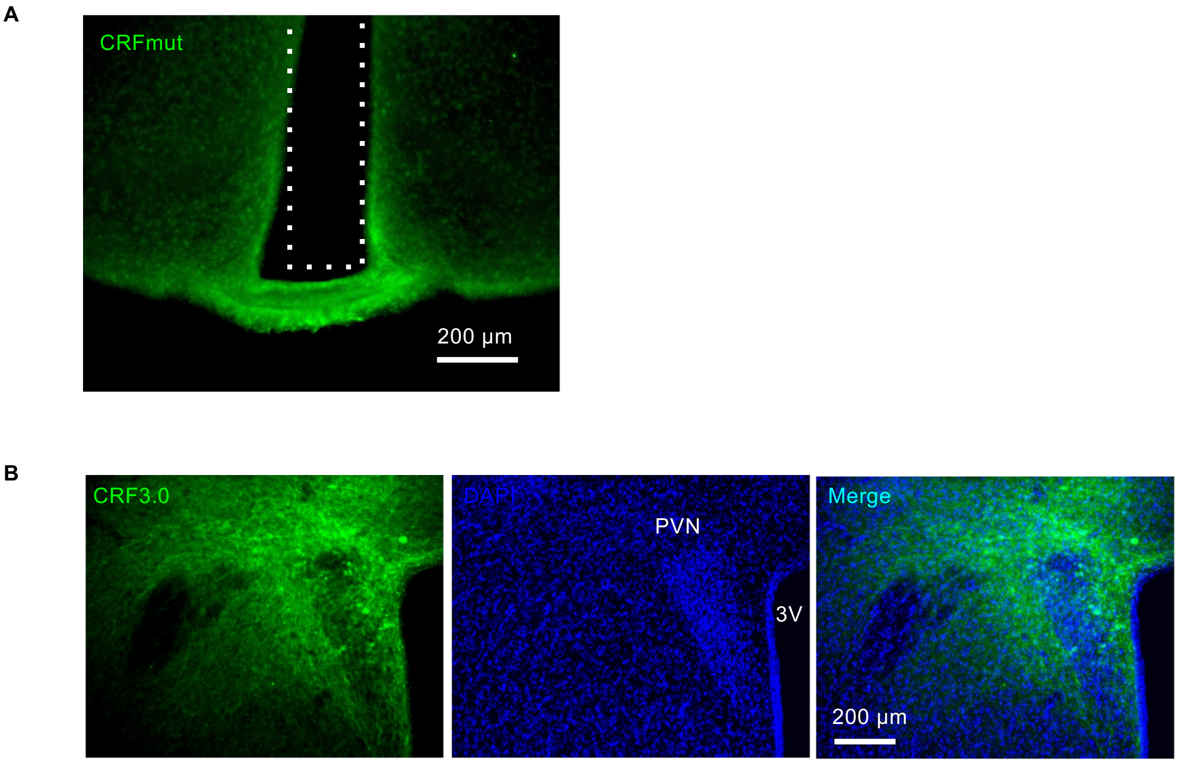


**Supplementary Figure 4. Photometry recording of CRH release at the ME.**
(A) The expression of CRH insensitive mutant sensor (CRFmut) at the ME. The optic fiber tract is marked with the dotted line in white. (B) The expression of CRF3.0 in the PVN (green) with the nuclear staining by DAPI (blue).

Supplementary Figure 5

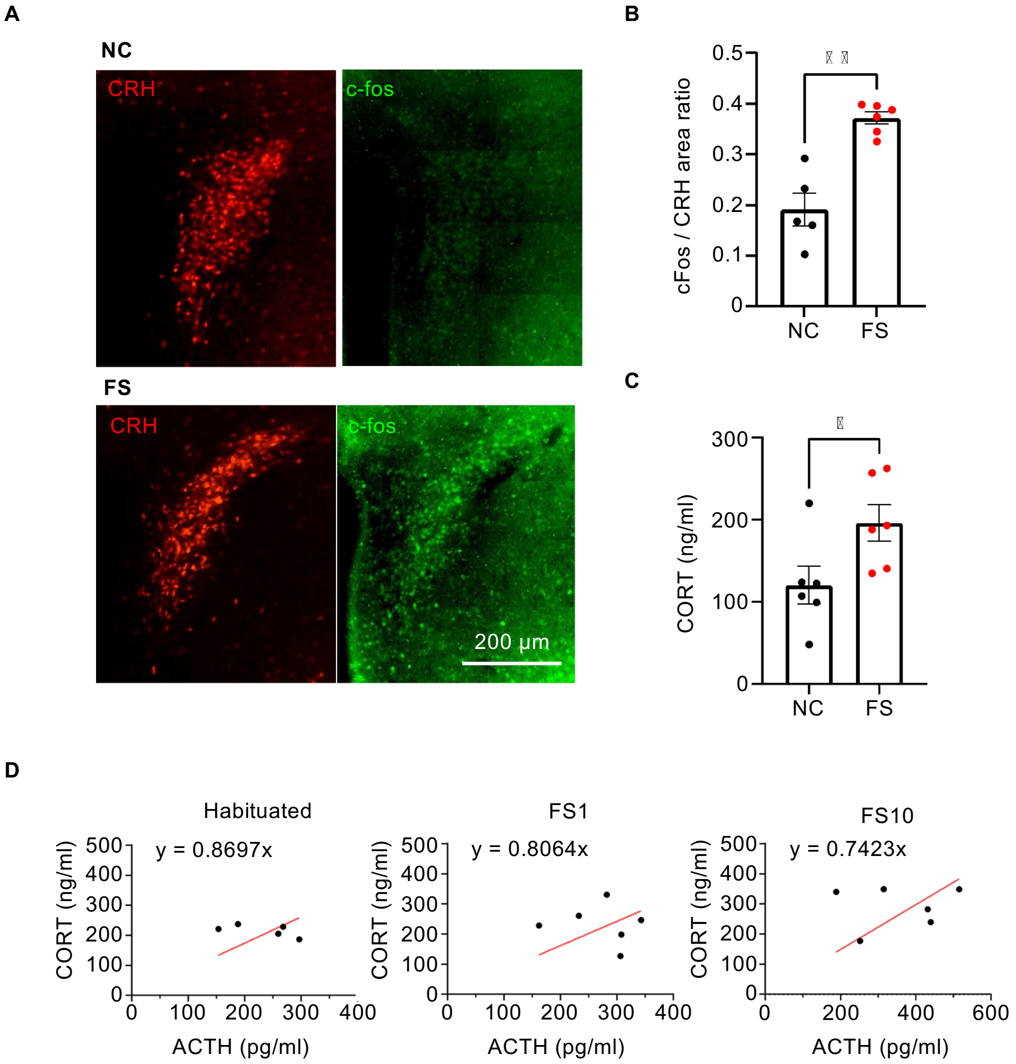


**Supplementary Figure 5. The activation of the HPA axis by foot shock stress.** (A) Representative images of CRH neurons (red) and c-fos-immunoreactivity (green) in the PVN after the foot shock. Top: negative control (NC) animal which were habituated to the chamber but not given any foot shocks. Bottom: Foot shocked animal which underwent 10 times foot shocks (FS10) . (B) Quantified data showing the ratio of total c-fos immunoreactive area divided by the total CRH positive area (**$p < 0.005$, Welch's *t*-test). (C) Plasma corticosterone (CORT) level of both NC and FS group (*$p < 0.05$, unpaired *t*-test). (D) Correlation between ACTH and CORT in the habituated(left), FS1(middle), and FS10(right) group. The equation of the fitting curve is shown on each graph.
